# *Staphylococcus aureus* wall teichoic acid is a pathogen-associated molecular pattern that is recognized by langerin (CD207) on skin Langerhans cells

**DOI:** 10.1101/238469

**Authors:** Rob van Dalen, Jacinto S. De La Cruz Diaz, Matevž Rumpret, Felix F. Fuchsberger, Nienke H. van Teijlingen, Jonas Hanske, Christoph Rademacher, Theunis B.H. Geijtenbeek, Jos A.G. van Strijp, Christopher Weidenmaier, Andreas Peschel, Daniel H. Kaplan, Nina M. van Sorge

## Abstract

*Staphylococcus aureus* is a major cause of skin and soft tissue infections and aggravator of the inflammatory skin disease atopic dermatitis (AD). Epicutaneous exposure to *S. aureus* induces Th17 responses through skin Langerhans cells (LCs), which paradoxically contribute to host defense but also to AD pathogenesis. The underlying molecular mechanisms of the association between *S. aureus* and skin inflammation are poorly understood. Here, we demonstrate that human LCs directly interact with *S. aureus* through the pattern-recognition receptor langerin (CD207). Human, but not mouse, langerin interacts with *S. aureus* through the conserved β-*N*-acetylglucosamine (GlcNAc) modifications on wall teichoic acid (WTA), thereby discriminating *S. aureus* from other staphylococcal species. Importantly, the specific *S. aureus* WTA glycoprofile strongly influences the level of Th1-and Th17-polarizing cytokines that are produced by *in vitro* generated LCs. Finally, in a murine epicutaneous infection model, *S. aureus* induced a more pronounced influx of inflammatory cells and pro-inflammatory cytokine transcripts in skin of human langerin transgenic mice compared to wild-type mice. Our findings provide molecular insight into the unique pro-inflammatory capacities of *S. aureus* in relation to inflammatory skin disease.

## Introduction

The inflammatory skin disease atopic dermatitis (AD, also known as eczema) affects up to 20% of children and 3% of adults worldwide (1). An important characteristic of AD is the disturbed microbiota composition with dominant presence of *Staphylococcus aureus*, but not of other staphylococcal species (2,3). In particular, the *S. aureus* CC1 lineage is overrepresented in AD isolates and was proposed to have particular yet-unidentified features that enable colonization of AD skin (4). Langerhans cells (LCs) are key sentinel cells in the skin epidermis and are implicated in *S. aureus*-induced skin inflammation. LCs are equipped with a diverse set of pattern-recognition receptors (PRRs) to sense intruders, including the LC-specific C-type lectin receptor (CLR) langerin (CD207) (5). LCs can phagocytose microbes and initiate adaptive immune responses by activating skin-resident immune memory cells or naïve immune cells in the lymph nodes (6,7). In response to *S. aureus*, LCs induce Th17 responses that help to contain *S. aureus* infection but paradoxically also aggravate AD (8,9). Despite the functional importance of LCs in *S. aureus*-mediated skin pathology, the molecular interaction between LCs and *S. aureus* and the functional response of LCs have received little attention.

A dominant and evolutionarily conserved component of the *S. aureus* surface is wall teichoic acid (WTA), which is important in nasal colonization, *S. aureus*-induced endocarditis, beta-lactam resistance and phage-mediated horizontal gene transfer (10-14). In the majority of *S. aureus* lineages, WTA is composed of 20-40 ribitol phosphate (RboP) repeating units modified with *D*-alanine and *N*-acetylglucosamine (GlcNAc). GlcNAc is linked to the anomeric C4 of RboP in either α or β configuration by glycosyltransferases TarM and TarS, respectively (12,15). Several *S. aureus* WTA glycoprofiles can be discriminated: WTA β-GlcNAcylation is conserved in almost all *S. aureus* strains, whereas WTA α-GlcNAcylation is only present in about one-third of the *S. aureus* isolates. A small selection of isolates even completely lack WTA glycosylation (10,16). Finally, WTA of *S. aureus* lineage ST395 is composed of a glycerol phosphate (GroP) backbone modified by *N*-acetylgalactosamine (GalNAc) (14). WTA glycosylation is an important determinant in host-pathogen interactions, which includes attachment to scavenger receptor SREC-1 in the nasal epithelium, and opsonization by antibodies and mannose-binding lectin (17-19).

We demonstrate an important role of the PRR langerin in sensing the β-GlcNAc epitope on *S. aureus* WTA, which explains the lack of binding to other non-AD associated staphylococcal species. Interestingly, simultaneous decoration of WTA with α-GlcNAc impairs langerin interaction and dampens cytokine responses of LCs, implying that *S. aureus* can modulate immune detection and subsequent inflammation in the epidermis. Murine infection experiments confirmed that langerin contributes to enhanced skin inflammation. In conclusion, we identify WTA as a pathogen-associated molecular pattern (PAMP) of *S. aureus*, which is recognized by langerin on LCs.

## Results

### Langerin is a receptor for *S. aureus* on human LCs

The molecular interaction between LCs and *S. aureus* has received little attention. We therefore investigated whether LCs and *S. aureus* interact directly by incubating primary LCs isolated from human skin with GFP-expressing *S. aureus*. LCs from four different donors bound *S. aureus* in a dose-dependent manner (Figure 1A). The levels at which the interaction was saturated varied between the donors from approximately 40% (donor 1) to 80% (donor 3) of *S. aureus*-positive LCs. To investigate the nature of interacting receptors on LCs, we pre-incubated LCs with mannan, a ligand for many PRRs of the CLR family. Depending on the bacteria-to-cell ratio, *S. aureus* binding was reduced by 35-70% compared to non-blocking conditions in all donors (Figure 1A). Similarly, the interaction was inhibited by approximately 35% by pre-incubation of the LCs with the monosaccharide GlcNAc (Figure 1A). Langerin is a mannan-and GlcNAc-specific CLR that is exclusively expressed on LCs. We therefore investigated whether langerin would be involved in interaction with *S. aureus*. Indeed, pre-incubation with an anti-langerin blocking antibody reduced binding of (*spa* and *sbi*-deficient, to prevent aspecific antibody binding) *S. aureus* in donors 3 and 4 by 25-50% compared to control, depending on the infective dose (Figure 1A). To confirm involvement of langerin in the interaction between *S. aureus* and LCs, we introduced langerin in the THP1 cell line, which normally does not express langerin. Transduction of langerin, but not of empty vector (EV), conferred *S. aureus* binding to THP1 cells, which could be completely inhibited by addition of mannan or anti-langerin blocking antibody (Figure 1B).

**Figure 1.**
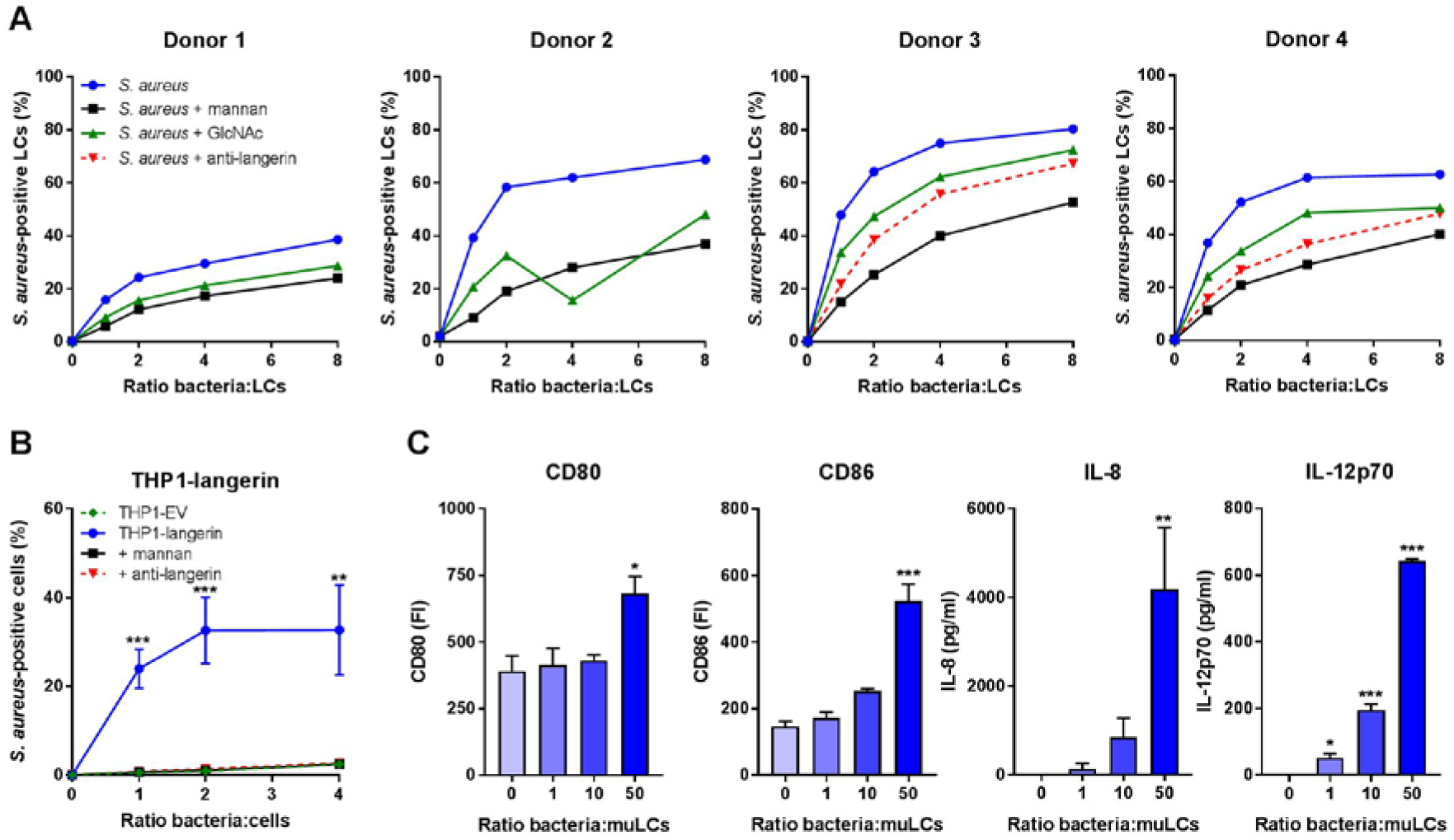
Langerin is a receptor for *S. aureus* on human LCs. (A) Binding of *S. aureus* to isolated primary human LCs. LCs from donors 1 and 2 were incubated with GFP-expressing *S. aureus* Newman and LCs from donors 3 and 4 with GFP-expressing *S. aureus* Newman Δ*spa*Δ*sbi* and binding was assessed by flow cytometry. The interaction was blocked by addition of mannan, GlcNAc or anti-langerin blocking antibody (donors 3 and 4 only). (B) Binding of *S. aureus* to THP1-langerin cells. Human langerin-transduced or empty vector (EV)-transduced THP1 cells were incubated with different amounts of GFP-expressing *S. aureus* Newman Δ*spa*Δ*sbi*. The interaction was blocked by addition of mannan or anti-langerin blocking antibody. Within each ratio, THP1-langerin was compared to the other conditions by two-way ANOVA followed by Dunnett’s multiple comparison test. (C) Expression of co-stimulatory molecules CD80 and CD86 and production of cytokines IL-8 and IL12p70 by muLCs after incubation with γ-irradiated *S. aureus* USA300 (24 h). Values were compared to unexposed control by one-way ANOVA followed by Dunnett’s multiple comparison test. Data are presented as percentage GFP+ cells (A, B), geometric mean fluorescent intensity or mean concentration (C) +/- standard error of mean (SEM) from three independent experiments, except for (A) (four donors with single measurements). \**P* < 0.05, \*\**P* < 0.01, \*\*\**P* < 0.001.

It was previously demonstrated that *S. aureus*-exposed LCs initiate T cell proliferation (20). However, the functional response of LCs was not assessed. Therefore, we stimulated MUTZ-3-derived LCs (muLCs), a well-established cell model for human LCs (21,22), with *S. aureus* and measured muLC activation through expression of co-stimulatory molecules and cytokine production after 24 hours. Indeed, muLCs upregulated expression of co-stimulatory molecules CD80 and CD86 and produced significant amounts of IL-8 and IL-12p70 in a dose-dependent response to *S. aureus* (Figure 1C). Together, these data demonstrate that LCs respond to *S. aureus* and that langerin is an important innate PRR for *S. aureus* on human LCs.

### Langerin recognizes *S. aureus* in a *tarS*-dependent manner through the conserved WTA β-GlcNAc epitope

To further investigate langerin interaction with staphylococci, we tested binding of a FITC-labeled trimeric construct of the extracellular domain of human langerin (langerin-FITC) to a broader collection of 18 *S. aureus* strains from 11 different clonal complexes, as well as several coagulase-negative staphylococci (CoNS). Langerin-FITC bound to most tested *S. aureus* strains but to none of the CoNS species (Figure 2A), indicating that langerin interacts with a ligand that is specific for and highly conserved in *S. aureus*. The three tested S*. aureus* strains that showed no or low-level binding of langerin-FITC (ED133, Lowenstein and PS187; Figure 2A), differ from the other tested *S. aureus* strains in the structural composition of WTA. ED133 and Lowenstein completely lack WTA GlcNAcylation, whereas PS187 belongs to the ST395 lineage that expresses GroP-GalNAc WTA (14,16,23). Given the high density of WTA on the *S. aureus* surface and apparent correlation between langerin interaction and WTA structure, we hypothesized that WTA GlcNAc modifications are likely candidates for the interaction with langerin.

**Figure 2.**
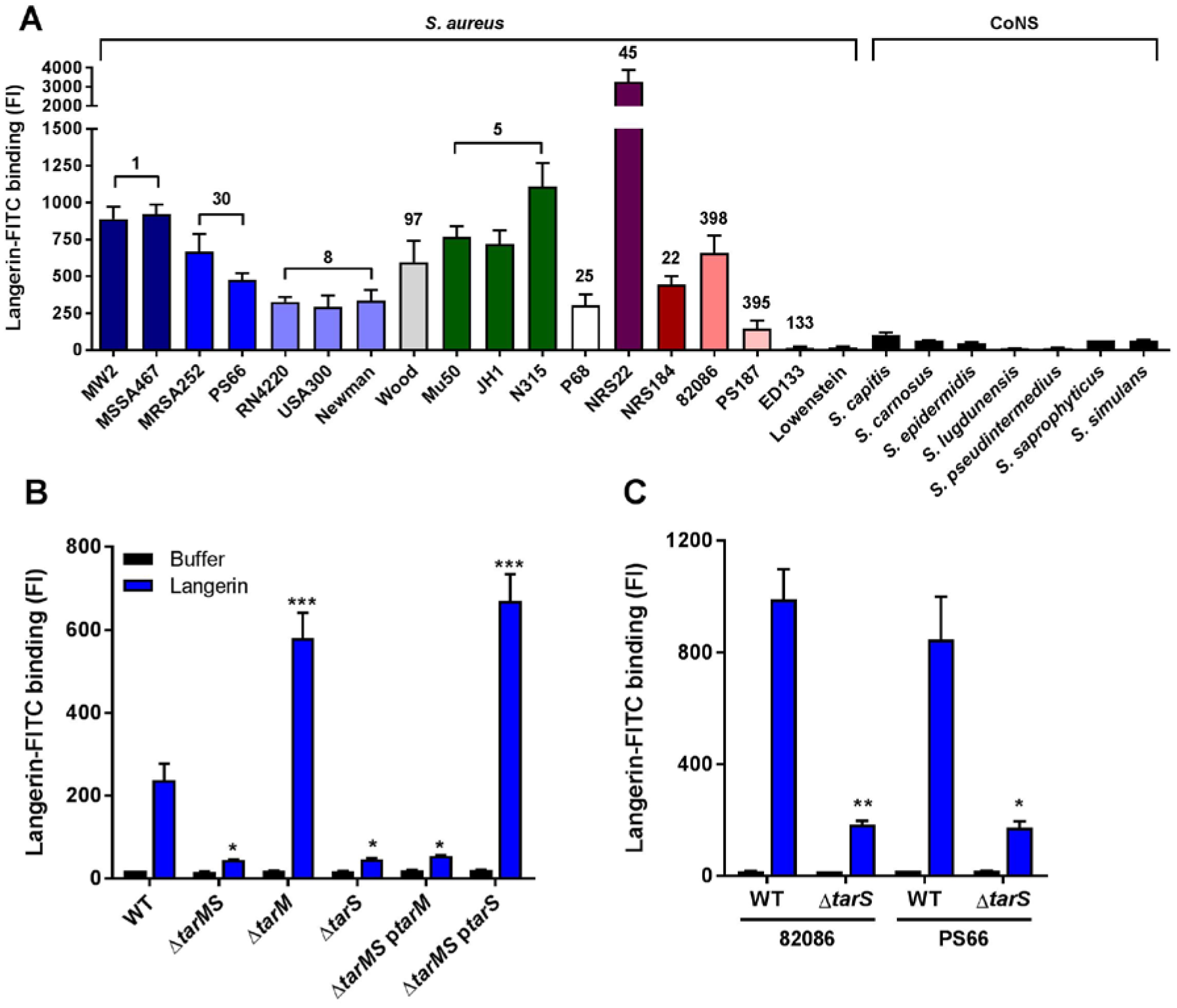
Langerin recognizes *S. aureus* in a *tarS*-dependent manner through the conserved WTA β-GlcNAc epitope. Binding of recombinant human langerin-FITC to (A) 18 wild-type *S. aureus* strains (11 different clonal complexes, indicated above the bars and by different color) and a selection of coagulase-negative staphylococcal species (CoNS); (B) *S. aureus* USA300 wild-type (WT) and WTA biosynthesis mutants Δ*tarMS*, Δ*tarM*, Δ*tarS*, Δ*tarMS* p*tarM* and Δ*tarMS* p*tarS*; and (C) two representative *S. aureus* isolates (82086 and PS66) that naturally lack *tarM* and their isogenic Δ*tarS* mutants. For B, C: all strains were grown to mid-exponential phase and incubated with langerin-FITC (blue) or buffer (black). Binding was assessed by flow cytometry. Data were compared by one-way ANOVA followed by Dunnett’s multiple comparison test (B) or by unpaired two-tailed *t*-test (C) and are presented as geometric mean fluorescence intensity + SEM from three independent experiments. \**P* < 0.05, \*\**P* < 0.01, \*\*\**P* < 0.001.

To test this hypothesis, we assessed binding of langerin-FITC to a panel of *S. aureus* knockout strains, which lack glycosyltransferases TarM and TarS that are required to modify WTA with α-GlcNAc and β-GlcNAc, respectively. Loss of both glycosyltransferases (Δ*tarMS*) reduced langerin-FITC binding to *S. aureus* to background levels in three different *S. aureus* backgrounds (Figures 2B and S1A, B), demonstrating that WTA GlcNAc is the target for langerin. To investigate whether langerin specifically recognized either α-GlcNAc or β-GlcNAc, we tested the individual TarM and TarS knockout strains as well as Δ*tarMS* complemented with either *tarM* or *tarS* on an expression plasmid (Δ*tarMS* p*tarM* and Δ*tarMS* p*tarS*). Langerin-FITC only bound to *S. aureus* strains that express β-GlcNAc, whereas α-GlcNAc was dispensable for binding (Figures 2B and S1A, B). Similarly, langerin-FITC binding to *S. aureus* strains 82086 and PS66, which are naturally deficient for WTA α-GlcNAc, was completely abrogated in isogenic Δ*tarS* strains (Figure 2C). These results show that langerin interacts with *S. aureus* in a *tarS*-dependent manner and provide the first demonstration of an anomeric-specific interaction of a human innate receptor with a Gram-positive surface polysaccharide.

Although α-GlcNAc is not the target of langerin, its presence influences the level of langerin-FITC binding: mutant strains lacking *tarM* (Δ*tarM* and Δ*tarMS* p*tarS*) showed significantly increased binding compared to wild-type (Figures 2B and S1A, B). Possibly, enhanced binding results from loss of steric hindrance by α-GlcNAc, since chemical analysis of the WTA composition by Kurokawa *et al*. suggests that WTA of strain RN4220 Δ*tarM* does not have increased β-GlcNAcylation (18).

As *S. aureus* expresses many human-specific adhesion or immune evasion factors (24), we investigated the interaction with murine langerin-FITC, which has 76% identity with the human langerin-FITC construct (25). Binding of murine langerin-FITC to *S. aureus* was detectable, but was 10 to 100-fold lower than human langerin (Figure S1C). The EC50 of human langerin-FITC for *S. aureus* USA300 was 9.7 (8.3 – 11.3) μg/ml, while binding of murine langerin-FITC was not yet saturated at 50 μg/ml. Despite low level and non-saturable binding, murine langerin interaction with *S. aureus* could be blocked by addition of mannan (data not shown), suggesting that the interaction is specific. Altogether, this indicates that the langerin-*S. aureus* interaction has a certain degree of species-specificity.

### *S. aureus* induces a Th1 and Th17 cytokine profile in LCs, which is affected by the WTA glycoprofile

Given the importance of langerin for interaction between *S. aureus* and LCs, we investigated whether distinct WTA GlcNAc glycoprofiles influenced the muLC response at the level of co-stimulatory molecules and cytokine expression. In line with our initial observations (Figure 1D), stimulation of muLCs with wild-type *S. aureus* upregulated expression of activation markers CD80, CD83 and CD86 (Figure 3A). Stimulation with β-GlcNAc-deficient *S. aureus* Δ*tarS* reduced expression of these markers compared to wild-type, whereas stimulation with α-GlcNAc deficient *S. aureus* Δ*tarM* enhanced expression (Figure 3A). In addition, muLCs secreted significant levels of cytokines IL-6, IL-8 IL-12p70, IL-23p19 and TNFα (Figure 3B), but not anti-inflammatory cytokine IL-10, in response to *S. aureus*. Cytokine levels were significantly reduced after stimulation of muLCs with *S. aureus* Δ*tarS* compared to WT, whereas stimulation with *S. aureus* Δ*tarM* significantly enhanced secretion of these cytokines (Figure 3B). Interestingly, the level of muLC activation correlated with the interaction levels of recombinant langerin-FITC to *S. aureus* WT, Δ*tarM* and Δ*tarS* strains (Figure 2B). These data suggest that the previously described Th1 and Th17-polarizing response initiated by LCs in response to *S. aureus* is strongly influenced by the specific glycoprofile of WTA.

**Figure 3.**
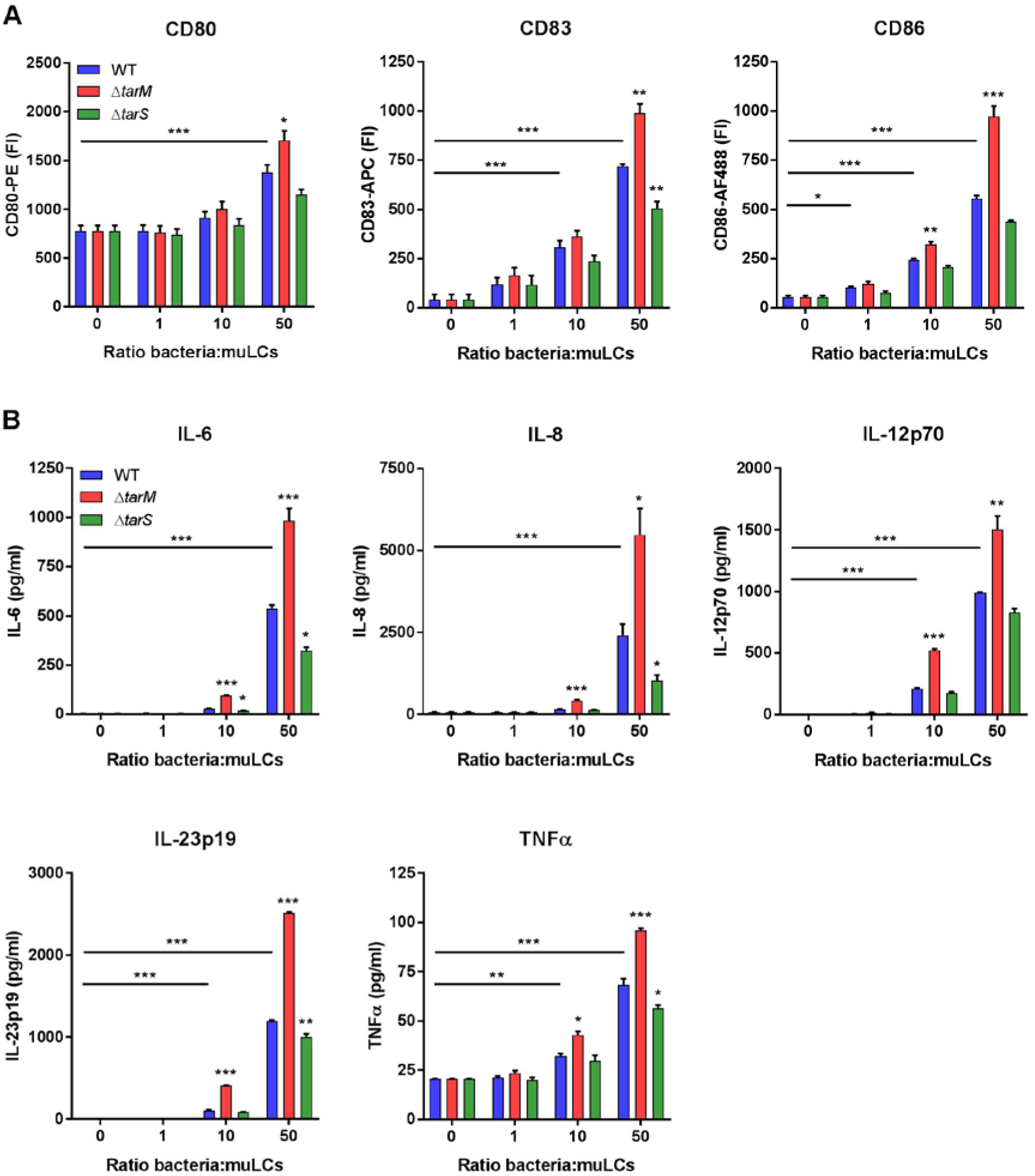
*S. aureus* induces a Th1 and Th17 cytokine profile in LCs, which is affected by the WTA glycoprofile. Expression of (A) co-stimulatory molecules CD80 and CD86 and maturation marker CD83 and (B) cytokines IL-6, IL-8, IL12p70, IL23p19 and TNFα by muLCs. muLCs were incubated with γ-irradiated *S. aureus* USA300 wild-type (WT), Δ*tarM* and Δ*tarS* for 24 h. muLCs stimulated with WT *S. aureus* were compared to the unstimulated control and muLCs stimulated with Δ*tarM* and Δ*tarS* were compared to their respective WT controls within the same ratio by one-way ANOVA followed by Dunnett’s multiple comparison test. Data are presented as geometric mean fluorescence intensity or mean concentration + SEM from three independent experiments. \**P* < 0.05, \*\**P* < 0.01, \*\*\**P* < 0.001.

### Human langerin transgenic mice show enhanced inflammation to epicutaneous *S. aureus* infection

Given the observed species specificity of langerin for *S. aureus* WTA β-GlcNAc (Figure S1A), we used human langerin - diphtheria toxin receptor (huLangerin-DTR) mice, which constitutively express human langerin on mouse LCs, as a huLangerin transgenic mouse model (26). Wild-type (WT) and huLangerin mice were epicutaneously inoculated with 1 x 10^7^ colony forming units (CFU) of *S. aureus* Δ*tarM* (Figure S2A) (8,27). These genetically stable mutant bacteria are unable to modulate WTA glycosylation through regulation of *tarM*, thereby maximizing the interaction with human langerin. At the time of sacrifice and skin collection (40 hours post-infection), the lesions of the huLangerin mice were clinically different from those of WT mice (Figure S2B), although bacterial burden in the skin did not differ between the groups (Figure S2C). Histological examination of the skin revealed more extensive influx of inflammatory cells in the dermis of huLangerin mice compared to WT mice (Figure 4A). Correspondingly, we observed significantly higher expression of the mouse IL-8 homolog CXCL1 (KC), but not of CXCL2 (MIP-2), in the huLangerin group as opposed to WT controls (Figures 4B and S2D). In addition, we determined the transcript levels of the Th17 cytokines IL-6 and IL-17 and the anti-inflammatory cytokine IL-10. Though not significant, both IL-6 and IL-17 showed a trend towards higher production in the huLangerin group, while IL-10 was not induced in either group, corroborating the observed *in vitro* responses of muLCs to *S. aureus* stimulation (Figures 4B and S2D). These results provide a first *in vivo* demonstration of the involvement of human langerin in the skin immune response to *S. aureus*.

**Figure 4.**
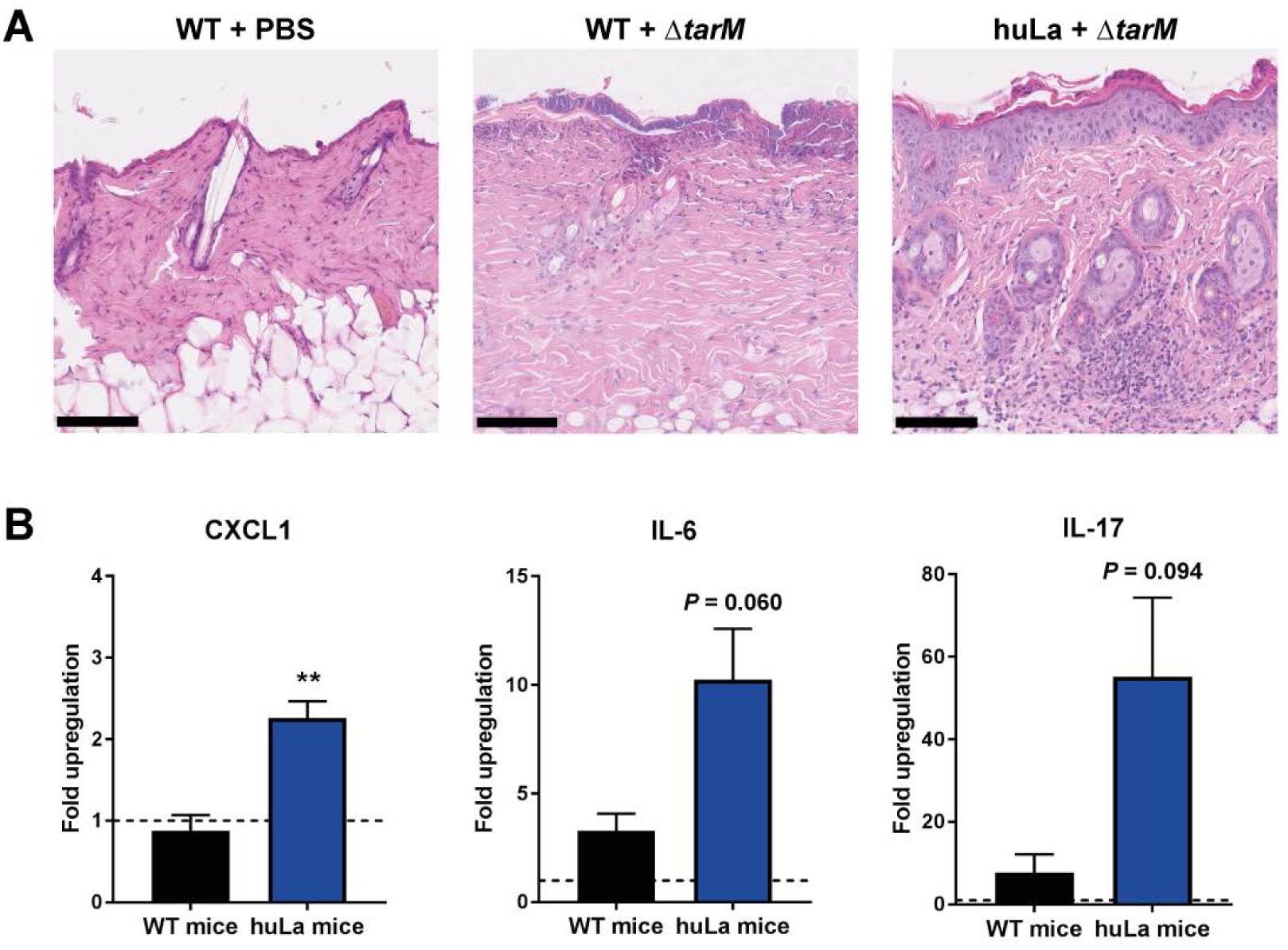
Human langerin transgenic mice show an enhanced inflammation to epicutaneous *S. aureus* infection. (A) Representative images of hematoxylin and eosin staining of skin biopsies and (B) transcript abundance of *CXCL1*, *IL-6* and *IL-17* from the lesions of WT (n=3) and huLangerin (n=4) mice 40 hours post-epicutaneous inoculation with *S. aureus* Δ*tarM*. The scale bars in (A) represent 100 μm. Data in (B) are presented as fold upregulation + SEM relative to *GAPDH* from three or four technical replicates and normalized for the WT/PBS control. The groups were compared by unpaired two-tailed *t*-test. \*\**P* < 0.01.

## Discussion

Despite the emerging role of LCs in *S. aureus*-mediated skin inflammation, there is limited information on the molecular pathways and functional consequences of LC - *S. aureus* interaction. We observe that LCs respond to *S. aureus* with a Th1 and Th17-polarizing cytokine response, which corroborates findings by others, who have demonstrated that LCs internalize *S. aureus* and subsequently polarize T cells towards Th17 (8,9,20,28). Furthermore, we elucidate that detection of *S. aureus* WTA β-GlcNAc is of critical importance for the induced cytokine response and can also be modified by co-decoration with α-GlcNAc, a characteristic of approximately one-third of *S. aureus* isolates. The ability of *S. aureus* to carefully regulate its WTA glycoprofile was also previously suggested in the context of lytic podophage infection (16). Likely, *tarM* is regulated as part of the GraRS regulon, known to control *S. aureus* susceptibility to antimicrobial defenses (29,30). However, whether and how GraRS and WTA GlcNAcylation are affected during skin colonization remains to be determined.

In addition to regulation of glycosylation, WTA abundance can be regulated through *tarH*, the ATPase required for WTA transport across the membrane (31). High WTA expression increases the ability to induce skin abscesses in mice (31). These results cannot be compared directly to our study, since the mice were infected subcutaneously, thereby bypassing the LCs. In addition, the species specificity of langerin should be taken into account. We demonstrate that mouse langerin shows significantly reduced binding to *S. aureus* compared to human langerin, underlining previous studies that reported differences in ligand specificity of these orthologs (25).

LCs and langerin were previously implicated in host defense against various other pathogens. LCs internalize and degrade HIV-1 viral particles in a langerin-dependent manner to prevent infection of deeper layers of the mucosa (32,33). Langerin has also been identified as a major receptor for fungal pathogens on LCs through recognition of mannose and beta-glucan structures (34). The Gram-negative bacterium *Yersinia pestis* is the only other bacterium known to interact with langerin and does so through its LPS core oligosaccharide (35). We identify *S. aureus* WTA β-GlcNAc as a new ligand for langerin. WTA is an abundant evolutionarily conserved feature of the surface of Gram-positive bacteria, making it advantageous for the host to recognize such structures in a timely manner. Although several receptors for *S. aureus* WTA have been described, langerin is the first human innate receptor to discriminate between the α-GlcNAc and β-GlcNAc modifications.

As an opportunistic microbial resident of the skin *S. aureus* is involved in the development of skin disease. Therefore, the recognition of *S. aureus* WTA by langerin on epidermal LCs that are strategically localized at mucosal surfaces may be key to maintaining skin homeostasis and preventing the development of infection or chronic inflammation. Alternatively, it is also possible that *S. aureus* exploits langerin interaction to intentionally elicit inflammation to perturb the skin barrier and release nutrients. Our epicutaneous infection experiments in WT and huLangerin transgenic mice do not support the latter hypothesis since we did not observe differences in bacterial burden despite increased inflammation in huLangerin transgenic mice. It remains to be determined why the clearly altered skin immune response in huLangerin mice does not result in differential bacterial survival in the skin. Potentially, 40 hours post-infection is too early to observe such an effect or the expression of immune evasion molecules allows survival of *S. aureus* in this hostile environment.

The identification of *S. aureus* as a new langerin-interacting pathogen is especially interesting in the context of AD. First, *S. aureus* is a driver of AD disease progression, which is mediated by LCs (9). Second, genome-wide association studies (GWAS) identified *CD207*, the gene encoding for langerin, as an AD susceptibility locus (36,37). In these studies, polymorphisms in a putative enhancer region of *CD207*, which likely increase expression of langerin, were protective for AD. Our data now functionally link langerin to *S. aureus*. Since *S. aureus* is largely resistant to host defenses but most of the other commensals are not, this could explain the strong association between *S. aureus* and AD, as well as the described driver function of *S. aureus* in AD disease progression. Also our observation that WTA α-GlcNAc attenuates LC activation can be important in the context of AD. The CC1 lineage is particularly overrepresented in isolates from AD skin and was suggested to have unidentified features that enable colonization by and proliferation of *S. aureus* on AD skin (4). Interestingly, all CC1 strains are *tarM*-positive (38), providing the potential to regulate WTA glycoprofile by co-decoration with α-GlcNAc. This could enable the bacteria to skew the inflammatory status of the skin and gain an advantage to colonize AD skin. Our data may provide molecular insight into the association between AD and *S. aureus* from two different angles: on the immunological side we show how langerin and LCs are involved in the immune response to *S. aureus*, while on the microbiological side the involvement of langerin could explain the association of *S. aureus* but not CoNS species with AD, and possibly also the overrepresentation of *tarM*-bearing CC1 strains in AD.

In conclusion, we identify *S. aureus* WTA as a PAMP and pinpoint langerin as a molecular trigger for *S. aureus*-induced skin immune responses. Our findings give a deeper understanding of the specific association of *S. aureus* with skin inflammation and can help in the development of new treatment strategies for *S. aureus*-associated skin and soft tissue infections and inflammatory skin diseases.

## Methods

### Bacterial strains and culture conditions

*S. aureus*, *S. capitis*, *S. carnosus*, *S. epidermidis*, *S. lugdunensis*, *S. pseudintermedius*, *S. saprophyticus* and *S. simulans* strains (Supplementary Table S1) were grown overnight at 37°C with agitation in 5 ml Todd-Hewitt broth (THB; Oxoid). For *S. aureus* strains that were plasmid complemented THB was supplemented with 10 μg/ml chloramphenicol (Sigma Aldrich). A fresh five ml THB culture was inoculated by 150 μl overnight culture and grown to an optical density at 600 nm (OD_600nm_) of 0.4 for *S. capitis* and to OD_600nm_=0.6-0.7 for all other bacteria, which corresponds to mid-exponential growth phase.

### Cell culture and muLC differentiation

MUTZ-3 cells (ACC-295, DSMZ) were cultured in a 12-well tissue culture plates (Corning) at a density of 0.5-1.0×10^6^ cells/ml in MEM-alpha (Gibco) with 20% fetal bovine serum (FBS, Hyclone, GE Healthcare), 1% GlutaMAX (Gibco), 10% conditioned supernatant from renal carcinoma cell line 5637 (ACC-35, DSMZ), 100 U/ml penicillin and 100 μg/ml streptomycin (Gibco) at 37°C with 5% CO_2_. We obtained MUTZ-3 derived Langerhans cells (muLCs) by differentiation of MUTZ-3 cells for 10 days in 100 ng/ml Granulocyte-Macrophage Colony Stimulating Factor (GM-CSF, GenWay Biotech), 10 ng/ml Transforming Growth Factor-beta (TGFβ; R&D Systems) and 2.5 ng/ml Tumor Necrosis Factor-alpha (TNFα; R&D Systems) as described previously (21,22). The phenotype of differentiated muLCs was verified by surface staining of CD34 (clone 581, BD Biosciences), CD1a (clone HI149, BD Biosciences) and CD207 (clone DCGM4, Beckman Coulter) using the respective antibodies and analysis by flow cytometry.

THP1 cells (TIB-202, ATCC) transduced with a lentiviral langerin construct or empty vector (EV) were cultured in RPMI (Lonza) supplemented with 5% FBS (Biowest), 1% GlutaMAX 100 U/ml penicillin and 100 μg/ml streptomycin (Gibco) at 37°C with 5% CO_2_.

### Isolation of primary human Langerhans cells

Human skin tissue was collected from otherwise healthy anonymous donors undergoing corrective breast or abdominal surgery. This study, including the tissue harvesting procedures, were approved by the Medical Ethics Review Committee of the Academic Medical Center Amsterdam, The Netherlands. Human Langerhans cells were isolated as described previously (33). In short, skin grafts were obtained using a dermatome (Zimmer) and incubated in medium supplemented with Dispase II (1 U/ml, Roche Diagnostics) after which epidermal sheets were separated from the dermis and cultured for three days. After incubation, migrated LCs were harvested and further purified using a Ficoll gradient (Axis-shield). Isolated LCs were routinely 90% pure (CD1a+ Langerin+) and were frozen in Iscoves Modified Dulbeccos’s Medium (IMDM, Thermo Fisher) supplemented with 20% FBS and 10% DMSO. Before use, LCs were thawed by drop-wise addition of cold IMDM with 10% FBS, washed twice and incubated in IMDM with FBS for 2 hours at 37°C with 5% CO_2_ to recover.

### Creation of GFP-expression *S. aureus*

To create GFP-expressing bacteria, *S. aureus* Newman wild-type and *S. aureus* Newman Δ*spa*Δ*sbi* were transformed as described previously with pCM29, which encodes superfolded green fluorescent protein (sGFP) driven by the *sarA*P1 promoter (39,40). In short, competent *S. aureus* were electroporated with pCM29 isolated from *E. coli* DC10B with a Gene Pulser II (BioRad; 100 Ohm, 25uF, 2.5kV). After recovery, bacteria were selected on TH agar supplemented with 10 μg/ml chloramphenicol. A single colony was grown in THB with 10 μg/ml chloramphenicol under the usual growth conditions. Bacterial expression of GFP was verified by confocal laser scanning microscopy (SP5, Leica).

### Gamma-irradiation of *S. aureus*

Gamma-irradiated stocks of *S. aureus* strains were made by harvesting cultures in mid-exponential growth phase by centrifugation (4,000 rpm, 8 min), which were concentrated 10x in phosphate-buffered saline (PBS; Lonza) with 17% glycerol (VWR), frozen at -70°C and exposed to 10 kGy of γ-radiation (Synergy Health, Ede, The Netherlands). Loss of viability of *S. aureus* was verified by plating of the irradiated bacteria. A non-irradiated aliquot that underwent the same freezing procedure was used to determine the concentration of colony forming units (CFU) of the irradiated stocks.

### Lentiviral transduction

A TrueORF sequence-validated cDNA clone of human CD207 (OriGene Technologies) was amplified by PCR using Phusion polymerase (Thermo Fisher) and primers hLangerin-Fw and hLangerin-FLAG-Rv (IDT, Supplementary Table S2). The PCR amplicon was cloned in a BIC-PGK-Zeo-T2a-mAmetrine;EF1A construct by Gibson assembly (NEB) according to the manufacturer’s instructions. The langerin-encoding vector and an empty vector (EV) control were introduced into THP1 cells by lentiviral transduction, as described by Van de Weijer *et al*. (41). In short, lentivirus was produced by HEK293T cells (CRL-3216, ATCC) in 24-well plates using standard lentiviral production protocols and third-generation packaging vectors. After 3-4 days the supernatant containing the viral particles was harvested and stored at - 70°C to kill any remaining cells. Approximately 50,000 THP1 cells were transduced by spin infection (1000xg, 2 h, 33°C) using 100 μl supernatant supplemented with 8 μg/ml polybrene (Santa Cruz Biotechnology). Complete medium was added after centrifugation and cells were selected three days post-infection by 100 μg/ml zeocin (Gibco). Cellular expression of langerin was verified by antibody staining of langerin (clone DCGM4, Beckman Coulter) and measured using flow cytometry.

### Bacterial binding assays

To test binding of bacteria to cells, 10^5^ LCs, THP1-EV or THP1-langerin were incubated with GFP-expressing *S. aureus* Newman or GFP-expressing *S. aureus* Newman Δ*spa* Δ*sbi* at bacteria-to-cell ratios from 1 to 8 in TSM buffer (2.4 g/L Tris (Roche), 8.77 g/L NaCl (Sigma Aldrich), 294 mg/L CaCl_2_·2H2O (Merck), 294 mg/L MgCl_2_·6H2O (Merck), pH=7.4) with 0.1% bovine serum albumin (BSA; Merck) for 30 minutes at 4°C. Binding was blocked by 15 minutes pre-incubation with 10 μg/ml mannan (Sigma Aldrich), 50 mM GlcNAc (Serva) or 20 μg/ml anti-langerin blocking antibody (clone 10E2, Sony Biotechnology). Cells were washed once with TSM 1% BSA, fixed in 1% formaldehyde (Brunschwig Chemie) in PBS and measured by flow cytometry.

### Production of recombinant langerin extracellular domains

The extracellular domains of truncated human langerin (residues 148–328) and mouse langerin (residues 150–331) were recombinantly expressed from codon-optimized constructs containing a C-terminal TEV cleavage site followed by a Strep-tag II cloned into pUC19 and pET30a (EMD Millipore) expression vectors as described previously (25). Recombinant human and murine ECDs were insolubly expressed in *E. coli* BL21(DE3), solubilized in 6 M guanidinium hydrochloride in 100 mM Tris (pH 8) with 1 mM DTT, refolded by dialyisis against Tris-buffered saline (pH 7.5) containing 10 mM CaCl_2_ and purified via mannan-coupled sepharose beads (Sigma Aldrich). Bound protein was eluted with Tris-buffered saline (pH 7.5) containing 5 mM EDTA. Protein concentrations were determined by A280 nm using the calculated molar extinction coefficients of 56,170 M^-1^ cm^-1^ for the human langerin ECD and 56,170 M^-1^ cm^-1^ for the murine ECD. The proteins were fluorescently labeled with fluorescein isothiocyanate (FITC, Thermo Fisher) by adding slowly 100 μL of the dye solution (1 mg/ml in DMSO) to 2 ml of a 2 mg/ml protein solution in HEPES-buffered saline (pH 7.2) containing 20 mM D-mannose (Sigma Aldrich) and 5 mM CaCl_2_. After stirring for 90 min at room temperature, the reaction was quenched by addition of 50 mM ethanolamine (pH 8.5, Sigma Aldrich). Unreacted dye molecules were removed by buffer exchange using a Zeba spin column (Thermo Fisher) and active protein was purified over mannan affinity column as described above. All chemicals used for the production of recombinant langerin extracellular domains were obtained from Carl Roth if not indicated otherwise.

### Langerin binding assay

Bacteria in mid-exponential growth phase were harvested by centrifugation (4,000 rpm, 8 minutes) and resuspended at OD_600nm_=0.4 in TSM buffer with 0.1% BSA. Bacteria were incubated with 1-50 μg/ml recombinant langerin-FITC (human or mouse) for 30 minutes at 37°C with agitation, washed once with TSM 1% BSA, fixed in 1% formaldehyde and analyzed by flow cytometry.

### muLC stimulation

We stimulated 5×10^4^ muLCs with *S. aureus* USA300 WT, USA300 Δ*tarM* or USA300 Δ*tarS* at bacteria-to-cell ratios of 0, 1, 10 and 50 in IMDM with 10% FBS. After 24 hours, supernatants were collected by centrifugation (300xg, 10 min, 4°C) and stored at -150°C until further analysis, and cells were washed once in PBS 0.1% BSA. Expression levels of the activation and maturation markers were determined by flow cytometry using the following antibodies: CD80 (clone 2D10), CD83 (clone HB15e) and CD86 (clone IT2.2, all from Sony Biotechnology) and their corresponding isotype controls (BD Biosciences).

### Cytokine assays

The IL-8 and IL12p70 concentrations were initially determined by ELISA (Sanquin and Thermo Fisher, respectively) according to the manufacturer’s instructions. Concentrations of IL-6, IL-8, IL-10, IL-12p70, IL-23p19 and TNFα cytokines were determined by Luminex xMAP assay (Luminex Corporation), performed by the Multiplex Core Facility UMC Utrecht, The Netherlands.

### Flow cytometry

Flow cytometry was performed on FACSVerse (BD Biosciences), per sample 10.000 events within the set gate were collected. Data were analyzed using FlowJo 10 (FlowJo, LLC).

### Epicutaneous murine infection model

We used 6-to 10 week-old sex-matched wild type C57BL/6 mice (obtained from Jackson laboratories) and huLangerin-DTR mice (26), provided by D. H. Kaplan (University of Pittsburgh, Pennsylvania USA). All mice were housed in a specific pathogen-free facility under standard conditions at the University of Pittsburgh. The mouse infection protocols were approved beforehand by the Institutional Animal Care and Use of Committee of the University of Pittsburgh. As described previously, mice were first anesthetized with a mixture of ketamine and xylazine (100/10 mg/kg body weight), shaved on the back with electric clippers, chemically depilated with Nair hair removal cream (Church & Dwight) according to the manufacturer’s instructions, and the stratum corneum was removed by 15 strokes of 220 grit sandpaper (3M) (8,27). After 24 hours, the mice were epicutaneously inoculated with PBS or *S. aureus* USA300 Δ*tarM*, which was grown overnight at 37°C in THB, in 50 μl of sterile PBS. Forty hours post-infection the mice were sacrificed and skin sections of 1 cm^2^ were collected. The sections were either 1) homogenized, serially diluted in sterile PBS, grown overnight on THB-agar plates at 37°C and colony forming units were counted, 2) homogenized and processed for RNA extraction or 3) fixed in 1% formalin in PBS. The fixed tissue sections were embedded in paraffin, cut, stained with hematoxylin and eosin, and digitalized (Hamamatsu NanoZoomer) by the Department of Pathology, UMC Utrecht, The Netherlands, and subsequently analyzed using NDP.view2.6.13 (Hamamatsu).

### Gene expression analysis

Whole skin was homogenized and processed for extraction and isolation of RNA, using TRIzol reagents (Thermo Fisher), following manufacturer’s instructions. RNA was quantified using a standard Nanodrop and cDNA was obtained using High-Capacity cDNA Reverse Transcriptase (Thermo Fisher). Quantitive PCR on cDNA was accomplished by using Taqman Gene Expression Master Mix and Taqman Gene Expression Assays for *IL-17*, *IL-6*, *CXCL1*, *CXCL2*, *IL-10* and *GAPDH* (Thermo Fisher) on a StepOnePlus Real Time PCR System (Applied Biosystems). Fold upregulation of transcripts was calculated from ΔΔCt values relative to *GAPDH* expression and normalized for PBS mock infection.

### Statistical analysis

Statistical analyses were performed using Graphpad Prism 7.02 (GraphPad Software). We used unpaired two-tailed *t*-tests for comparisons between two groups and one-way ANOVAs with a common control group followed by Dunnett’s multiple comparisons test. The THP1-langerin dose-response curves were tested using a two-way ANOVA followed by Dunnett’s multiple comparison test and the langerin-FITC concentration curves were tested against wild-type langerin-FITC using a two-way ANOVA followed by Tukey’s multiple comparison test. Data are presented as the geometric mean or percentage positive cells (flow cytometry), mean concentration (cytokine arrays) or fold upregulation (real-time PCR) + standard error of the mean (SEM).

### Data availability

The data that support these findings are available from the corresponding author upon request.

## Acknowledgements

The authors would like to thank: the Multiplex Core Facility (UMC Utrecht, The Netherlands) for performing the Luminex assay; David Gerlach, Guoqing Xia and Volker Winstel (currently or previously at University of Tübingen, Germany) for advice and communication regarding the shipment of strains; Samantha van der Beek, Eline van Yperen and Małgorzata Mnich (UMC Utrecht, The Netherlands) for technical assistance; Tanja de Gruijl (VU University Medical Center, Amsterdam, The Netherlands) for the protocol detailing MUTZ-3 culture and differentiation. This work was supported by a VIDI grant (91713303) from the Dutch Scientific Organization (NWO) to N.M.v.S and R.v.D. A.P. is supported by German Research Council (DFG) grants TRR34, TRR156 and the German Center for Infection Research (DZIF). J.H. thanks the DFG Collaborative Research Centre 765 “Multivalency” for a fellowship. C.R. thanks the DFG for funding (RA1944/2-1).

## Author Contributions

R.v.D., M.R. and N.M.v.S. planned the experiments. R.v.D., J.S.D.L.C.D. and M.R. performed the experiments and prepared the figures. F.F.F., J.H. and C.R. supplied the langerin-FITC constructs, N.H.v.T. and T.B.H.G. provided the primary LCs, C.W. and A.P. provided the bacterial strains, D.H.K. provided the mice. R.v.D. and N.M.v.S. wrote the manuscript, N.H.v.T., J.A.G.v.S., C.W. and A.P provided critical feedback.

